# CaviDB: a database of cavities and their features in the structural and conformational space of proteins

**DOI:** 10.1101/2022.08.02.502569

**Authors:** Ana Julia Velez Rueda, Franco Leonardo Bulgarelli, Nicolás Palopoli, Gustavo Parisi

## Abstract

Proteins are the functional and evolutionary units of cells. On their surface, proteins are sculpted into numerous concavities and bulges, offering unique microenvironments for ligand binding or catalysis. The dynamics, size, and chemical features of these cavities are essential for the mechanistic understanding of protein function.

Here we present CaviDB (https://www.cavidb.org), a novel database of cavities and their features in known protein structures, which integrates the results from commonly used software for cavities detection with protein features obtained from sequence, structure, and function analyses. Additionally, each protein in CaviDB is associated with its corresponding conformers which help to analyze conformational changes in cavities as well. We were able to characterize a total number of 16,533,339 cavities, 62,0431 of them predicted as druggable targets. CaviDB contains 276,432 different proteins, with information about all their conformers. It also offers the capability to compare cavities and their features from different conformational states of the protein. Furthermore, we have recently added the available models from the AlphaFold database versions 2 and 3, which allow further cavity explorations and comparisons.

Each entry information is organized in sections, highlighting the general cavities descriptors, including the inter-cavities contacts, activated residues per cavity, the information about druggable cavities, and the global protein descriptors. The data retrieved by the user can be downloaded in a format that is easy to parse and integrate with custom pipelines for protein analysis.

CaviDB aims to offer a comprehensive database for use not only in different aspects of drug design and discovery but also to better understand the basis of the protein structure-function relationship better. With its unique approach, CaviDB provides an essential resource for the wide community of bioinformaticians in particular and biologists in general.

## Introduction

Proteins are the functional, structural, and evolutionary units of cells. They are composed of chains of amino acids interacting in complex and highly connected networks. On their surface, proteins are sculpted into numerous concavities and bulges, offering unique microenvironments for ligand binding or catalysis [15]. The dynamics of these cavities are fundamental for understanding protein function and their variations can explain changes in protein activity [20,46]. Protein movements, even the smallest ones, can have an impact on the architecture of the cavities [18,25]. On different time scales, the movements are required not only to bind the substrate or to define its affinity constant but also to allow ligand transit from the surface to the active site [38].

The size and geometry of the cavities, as well as their accessibility, have proven useful in making predictions about protein-protein interactions, protein pharmacology, and binding specificity [5,8,30]. For instance, physicochemical properties of the cavities such as their charge or hydrophobicity can also be used to predict the binding probability of certain ligands [1,63]. Indeed, it is well established that the residues can shift their pKa values due to different structural and environmental features [3,14], favoring several biological activities [17,19]. Furthermore, it was shown that the shape and location of other cavities close to each other can define their relative flexibility and affect their catalytic and binding promiscuity [5,45,53].

Functional cavities are usually found within protein domains, that is, evolutionarily conserved protein regions with a certain stability, function, and dynamics. Strictly speaking, domains are a portion of the protein that may have none, one or more functional cavities. The biological roles of singular cavities and domains may not always be correlated and the conservation of cavities may exceed that of a certain domain family. Thus, knowing the activity of the domains is insufficient to fully understand the specific function of the proteins, and an integrative characterization of the complete protein and its cavities may be a better approach to knowing the protein function [50].

Here we present CaviDB (http://www.cavidb.org/), a novel database that integrates the results from commonly used software for cavities detection with protein features obtained from the sequence, structure, and function analyses, presented in an interactive online database. In particular, CaviDB integrates well-established methods for cavities detection [49,67] that enables the local structural characterization but is also helpful to understand the anatomy and function of the protein at a global level [9]. Our database also allows users to explore the dynamics of protein through an easy-to-use interface for comparing the features of protein conformers and their predicted cavities. CaviDB yields structural data about every protein structure known and available in the Protein Data Bank [55] and the predicted proteomes available for common study organisms in the AlphaFold database [24]. We aim to offer a comprehensive resource for use in different biotechnological application fields, such as drug design and discovery, but also to better understand the basis of protein structure-function relation.

## Materials and Methods

### Cavities prediction and categorization

CaviDB provides users with structural and sequential features for protein cavities characterization. The cavities predictions were made running the widely adopted software Fpocket [31] with its default settings, for all the Protein Data Bank [4,41], and all the Alphafold database entries [58]. We retrieved and annotated all the properties linked to each cavity and to all cavity-lining residues. The cavity was considered druggable if it displayed a druggability score greater than 0.5, as proposed in previous works [49].

### Cavities features calculation

In order to provide users with information about possible activated cavities, we estimated the pKa values (at pH=7) of the ionizable residues and their shifts (pKa predicted - pKa expected) using PROPKA [40]. The net pKa shift values per cavity were calculated as the sum of all absolute pKa shifts of each ionizable residue belonging to a cavity.

We also retrieved data about interresidue contacts per site using PROPKA, in order to annotate the cavities’ contacts as side chain hydrogen bonds, and backbone hydrogen bonds) and coulombic bonds. We built a network of cavities that share at least one contact between the same sites, which can be inspected as an interactive plot. The binding energies matrices were built by calculating the sum of absolute binding energies between the residues in contact in each pair of cavities.

The properties per site were calculated using CIDER [21], modlAMP [37], and Biopython [6], and assigned to each cavity as the mean values of the features of its residues.

### Global protein features calculation and annotation

The global protein features were calculated as described in the previous section. Each PDB or AlphaFold model was annotated through SIFTS [59] with identifiers of relevant biological databases, such as CATH [51] and Pfam [13], in order to help users with posterior analyses.

### Conformation comparison

For the conformers’ cavities comparisons, we used the PDBSWS - PDB/UniProt Mapping [32]. This database maps PDB residues to residues in UniProtKB (SwissProt and trEMBL) entries, consequently allowing the precise comparison between cavities of different entries.

### Web application overview

A responsive web interface was built to show the data stored in a non-relational database, which supports more straightforward navigation and visualization of the database contents on multiple devices. The web application was implemented in HTML, CSS, Ruby (on Rails), and JavaScript (with NodeJS).

The first step for running CaviDB is providing a valid PDB or Uniprot ID. The web server will automatically load all the chains related to the search and their general data including their length, the number of cavities predicted, and relevant cross-reference identifiers (Fig. 1A.). The search can be filtered with the AlphaFold selector when interested in these entries only. The features obtained for each entry are organized in two main sections describing the general cavities descriptors, including an interactive display for cavities visualization, a network plot of interactions, and the cavities including activated residues with pKa shifts, and the global protein descriptors (Fig. 1D.).

**Fig.1.**
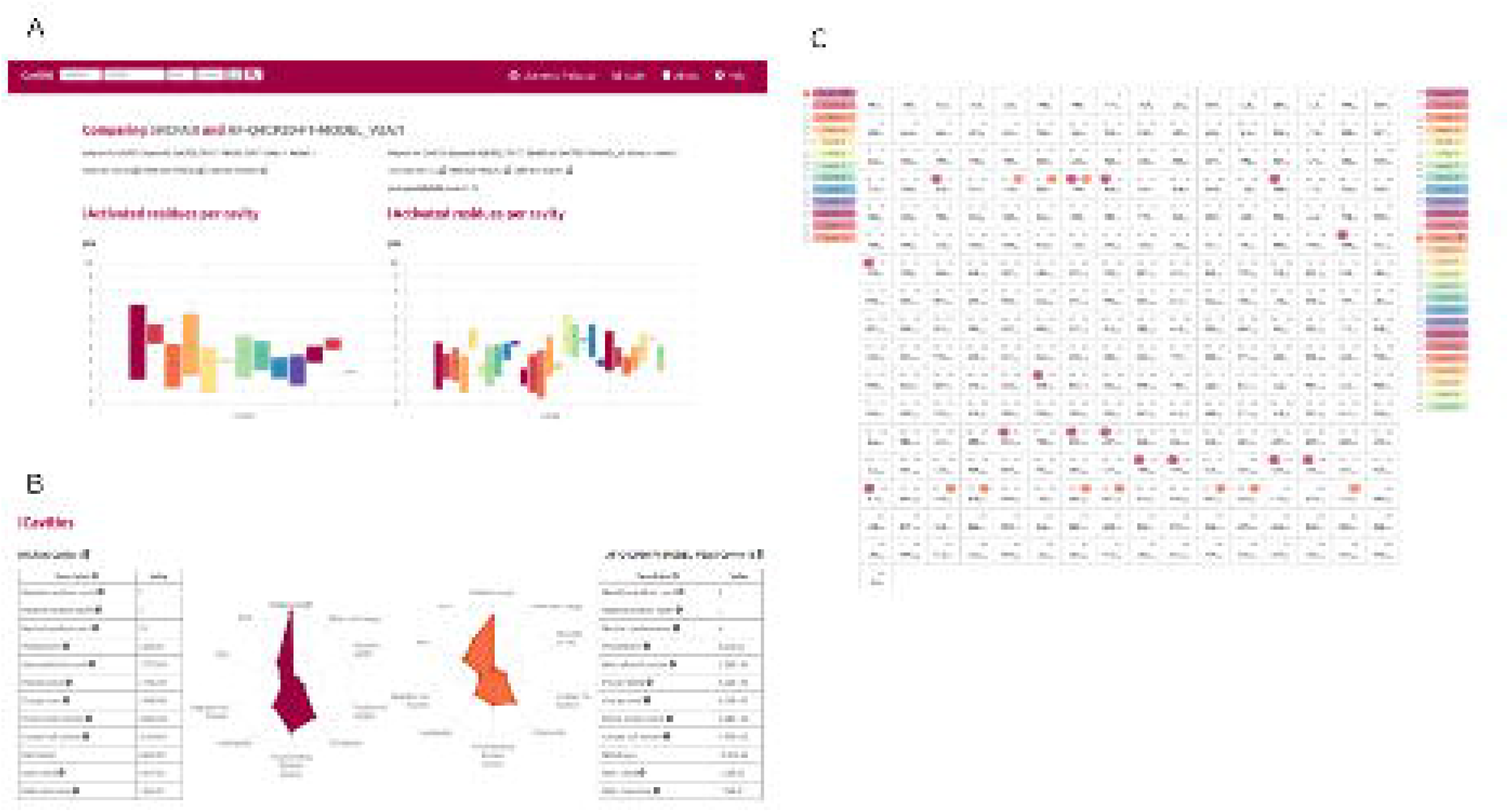
CaviDB web application overview: A) CaviDB search allows users to search by a specific PDB or Uniprot identifier. Also, a selector for focusing the search on AlphaFold models is provided; B and C) The cavities dynamics can be explored by using the conformation comparator provided by CaviDB, in which the predicted cavities and their features for different protein conformations could accede; D) Chain features’ renderization schematic example: each entry’s information is organized in two main sections, one containing the general cavities descriptors, and the global protein descriptors.

CaviDB allows users to explore the conformational diversity of proteins and their impact on the cavities dynamics by providing a conformation comparator (Fig. 1B.), that renders a comparison page with the listed cavities for each chain and, when selected, their characteristics and residues.

### Globular Protein Test Case

Promiscuous proteins are a breaking point in the ‘structure–function’ paradigm and the biological specificity concept [2,26]. The promiscuous behavior of proteins offers both challenges and opportunities to drug discovery programs and has been explored as a strategy for drug repurposing [10,16,57].

Human Serum Albumin (HSA) is the main protein in plasma, binds multiple ligands [60], and has recently emerged as a very important drug carrier [33,65]. HSA has been previously described not only as a transport protein but also as a promiscuous enzyme possibly linked to salicylic acid metabolism and with side effects [45,48,52,62,64].

It was proposed that the basis of the great capability of albumins to catalyze different reactions relays in the existence of activated amino acids, with abnormal pKas [22,45] in the hydrophobic cavity of the AII binding site, which creates a microenvironment favorable for catalysis. Indeed, as shown in the solvent accessibility (ASA) per site plot built by CaviDB for the 1AO6A:0 entry, there is a local minimum in the surrounding area of Lys199 (see Fig. 2B.), a region described as important for catalysis [29,64]. CaviDB identifies the AII binding site as the largest cavity (I), identified with the highest relative parameter (equal 1), showing a large number of inter-cavity contacts and the presence of activating residues. A second druggable cavity (II) is also identified which contains the Arg410 and Tyr411 residues that were previously described as part of the catalytic active site [62] (Fig. 2.). In addition, Tyrosine 411 and Arginine 410 (cavity 4), two residues shown to be important for the protein esterase-like activity [29], interact through coulombic forces (Fig. 2B.).

**Fig.2.**
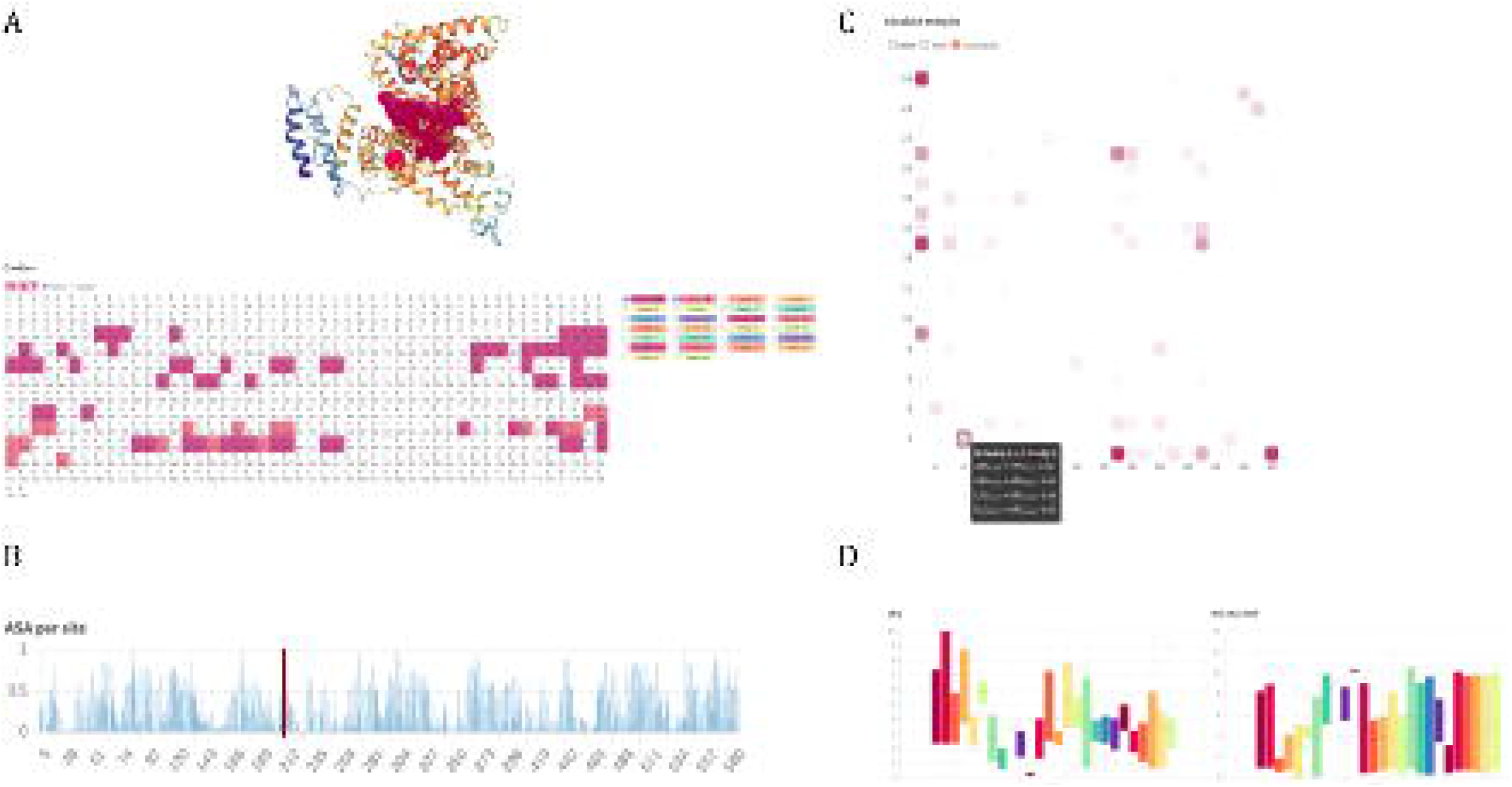
Human Serum Albumin (PDB ID: 1AO6, chain: A, model: 0) in CaviDB. A) Protein display and residues per cavity, with cavity I and II highlighted in the sequence (magenta and pink, respectively); B) Global accessibility per site plot: in red residue the surrounding of Lys199, in a local minimum ASA; C) Network plot of interactivities contacts, showing the 2 druggable cavities highlighted with a black border, above a heatmap display of all possible pairwise contacts. The Tyr411 and Arg410 interactions are highlighted in the box; D) Boxplot distributions of pKa values (left) and pKa shifts (right) per residue in each detected cavity.

### Intrinsically Disordered Protein Test Cases

In recent years, research about intrinsically disordered proteins (IDPs) as important targets for drug design has been growing [7,23,34]. In this sense, CaviDB is enriched with the three-dimensional structures predictions of proteins available in the Alphafold database, to complement the information on binding sites and cavities for which there is no crystallographic information [47,54].

An interesting case of study of ID proteins is the Calreticulin (CRT) protein from *T. cruzi*. This protein plays a very important role in parasite-host interactions and in the host’s tumor growth [11,44]. Upon infection, the parasite exposes the protein on its surface and promotes parasite infectivity [12,35]. CRT exhibits 3 disordered regions reaching 25% [42], and a global fraction of disorder-promoting residues [56] of 6.88E-01 calculated by CaviDB. CRT has 7 conformations stored in CaviDB, 6 coming from PDB, and 1 from AlphaFold. It presents a P-domain with a high-affinity calcium-binding site (from residue 205 to 302) [36], that has been shown to be involved in protein-protein interaction. Even when the amino acid sequence of the P domain is highly conserved, it includes flexible regions. A straightforward comparison between conformations can be done by using CaviDB. A quick inspection of the comparison between 5HCFA:0 and AF-Q4CPZ0-F1-MODEL_V2A:1 (pLDDT: 92.15) shows the great difference in the length of the chains due to the disorder regions, and therefore in the number of cavities predicted for one and the other conformation. When closely checking the activated cavities shown in CaviDB, it is interesting to notice that cavity 4, which includes the Cl binding residues (60 and 62) has a median pKa higher than 5 (Fig.3A.). The carbohydrate-binding sites (108,126,133, and 316) are part of the activated cavities 5, 17, and 14 include (Fig. 3A.). Finally, the p-domain that is missing in the PDB [36] and is modeled by AlphaFold, also has cavities (20,22 and 23) lying in this region.

**Fig.3.**
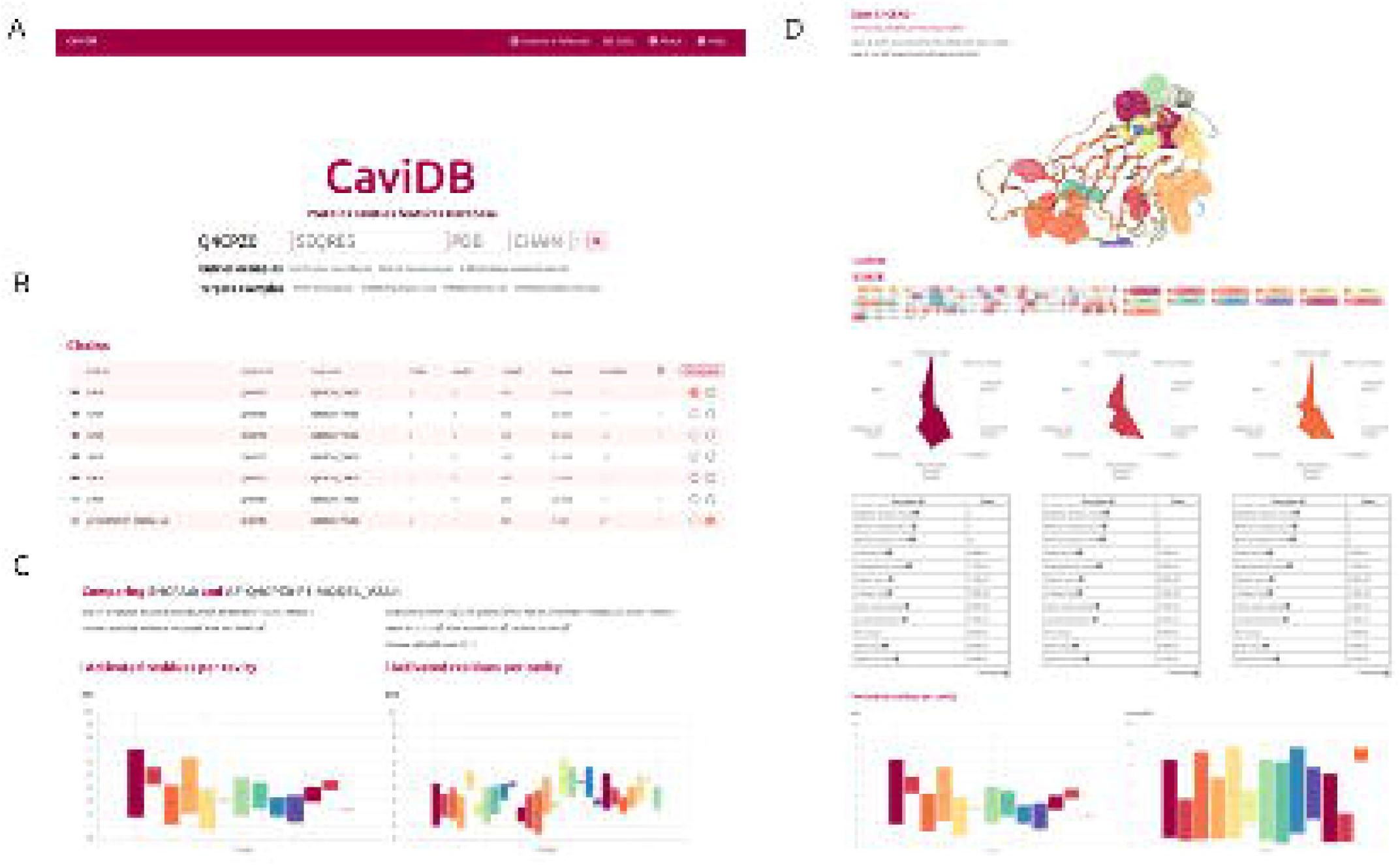
Comparison of Calreticulin (CRT) protein conformations. A) General conformers description and activated cavities data for AF-Q4CPZ0-F1-MODEL_V2A:1 and 5HCFA:0 entries; B) Druggable cavities 13 in AF-Q4CPZ0-F1-MODEL_V2A:1 chain which is the only druggable cavity detected, and partially overlaps with cavity 1 in 5HCFA:0, but shows a larger volume and polarity, and a lower mean pKa (Fig. 3.); C).

### The advantages of CaviDB over existing services

CaviDB has a total number of 276,432 different proteins, with 716,260 conformers coming from PDB and 278,420 from the AlphaFold Database. We have proteins from 7,191 species, representing a total number of 9,827 Pfam families and 415,173 Cath superfamilies. With the current number of entries, we were able to characterize a total number of 16,533,339 cavities, 62,0431 of them druggable. CaviDB can be found the information about 3,706 different enzymes, with 1,012 entries corresponding to Oxidoreductases class (EC1), 1,170 Transferases (EC2), 1,023 Hydrolases (EC3), 497 Lyases (EC4), 238 Isomerases (EC5) and 173 entries corresponding to Ligases (EC6). Since CaviDB provides the gene ids and Ensembl ID, each entry data could be easily mapped with metabolic pathways data and evolutionary information in which each protein may be involved.

There are many tools oriented to protein structural characterization and cavities prediction [28,61], and the number of 3D structures data is growing in big steps [24]. Nevertheless, as we have shown CaviDB is not only a useful tool for obtaining protein cavities features and their dynamics but also provides an easy and accessible way of analyzing structural data.

## Discussion

Identifying binding cavities is essential for understanding the relationship between structure and function in proteins and an essential step for drug design [39,61,63,66]. However, over the last few years, the number of three-dimensional protein structures has increased considerably [27,43]. But the more the data becomes available, the more challenging it’s structuration and processing. In this regard, CaviDB not only provides a freely accessible, comprehensive protein and their cavities features database, but also a simple and user-friendly tool for analyzing the data with a dynamic perspective, at multiple levels.

Our database also provides general annotations for the cavity-containing proteins, such as cross-references with CATH and Pfam [13,51], which enables further analysis of protein functionality based on the structural information.

## Author contributions

AJVR conceived the study and was in charge of the overall planning and direction of the project. GP and NP were in charge of the supervision and theoretical validation. AJVR and FB carried out the software development and implementation. FB made the technological supervision. AJVR, GP, NP, and FB wrote the manuscript.

## Declaration of Competing Interest

The authors declare that they have no known competing financial interests or personal relationships that could have appeared to influence the work reported in this paper.

## Funding

AJVR is a Postdoctoral fellow from CONICET. GP and NP are researchers from CONICET. This work was supported by Universidad Nacional de Quilmes (PUNQ 1004/11), ANPCyT (PICT-2014-3430, PICT-2013-0232), and AWS-CONICET INNOVA 2021 (Project 2022011357003403). The funders had no role in the study design, data collection, and analysis, decision to publish, or preparation of the manuscript.

## Notes

https://www.cavidb.org

### Competing Interest Statement

The authors have declared no competing interest.

